# Males that silence their father’s genes: genomic imprinting of a complete haploid genome

**DOI:** 10.1101/2020.04.27.063396

**Authors:** Andrés G. de la Filia, Andrew J. Mongue, Jennifer Dorrens, Hannah Lemon, Dominik R. Laetsch, Laura Ross

## Abstract

Genetic conflict is considered a key driver in the evolution of new reproductive and sex determining systems. In particular, reproductive strategies with non-Mendelian inheritance, where parents do not contribute equally to the genetic makeup of their offspring. One of the most extraordinary examples of non-Mendelian inheritance is paternal genome elimination (PGE), a form of haplodiploidy which has evolved repeatedly across arthropods. Under PGE, males are diploid but only transmit maternally-inherited chromosomes to their offspring, while the paternal homologues are excluded from sperm. This asymmetric inheritance is thought to have evolved through an evolutionary arms race between paternal and maternal genomes over transmission to future generations. In several clades with PGE, such as the mealybugs (Hemiptera: Pseudococcidae), paternal chromosomes are not just eliminated from sperm, but also heterochromatinised early in development and thought to remain inactive. Such paternal genome silencing could alleviate genetic conflict between paternal alleles over transmission. However, it is unclear if paternal chromosomes are indeed genetically inert in both soma and germline. Here, we present a parent-of-origin allele-specific transcriptome analysis in male mealybugs. We show that expression is globally biased towards the maternal genome, but detect activity of paternal chromosomes in both somatic and reproductive tissues. Up to 70% of somatically-expressed genes are to some degree paternally-expressed. However, paternal genome expression is much more restricted in the testis, with only 20% of genes showing paternal contribution. Finally, we show that the patterns of parent-of-origin-specific gene expression are remarkably similar across genotypes and that those genes with biparental expression show elevated rates of molecular evolution. Our results provide the clearest example yet of genome-wide genomic imprinting (parent-of-origin specific gene expression) in insects. Furthermore, it enhances our understanding of PGE, which will aid future empirical tests of evolutionary theory regarding the origin of this unusual reproductive strategy.

## Introduction

Sex—the mixing of heritable material of two individuals—is nearly universal among multicellular organisms. Yet the variety of ways organisms achieve this is staggering. For example, organisms can differ in the way genetic material is inherited, how the genes of two parents are combined to form offspring, or how offspring sex is determined. Why there is such large variability in a process that is so fundamental remains one of the unsolved mysteries of life. A rapidly growing body of theoretical literature suggests that conflicts between the sexes play a central role in driving the rapid turnover of reproductive and sex determining systems (Hurst et al., 1996; van Doorn and Kirkpatrick, 2007, 2010; Bachtrog et al., 2014; Gardner and Ross, 2014; Úbeda et al., 2015). These conflicts are particularly pronounced during reproduction, as they force unrelated mates to cooperate, and whilst this cooperation is aimed at achieving a common goal—producing descendants—not all interests are aligned (Chapman et al., 2003).

One particular conflict between parents that could have led to the evolution of new reproductive strategies is over the inheritance of their genes to their offspring (intragenomic conflict between parents: Hurst, 1992; Haig, 2000; Burt and Trivers, 2006; Ross et al., 2010; Werren, 2011; Normark and Ross, 2014; Gardner and Úbeda, 2017). Despite these conflicts, in many species—like ourselves—reproduction is a fair process that follows Mendel’s classic laws of inheritance: each parent contributes an equal share of the heritable material of their offspring (Wright, 1931). Yet in some species (approximately 15% of animals), these rules of fair reproduction are violated (Bachtrog et al., 2014). Probably the best-known example is haplodiploidy—found in wasps, ants and bees—where sons only inherit their mother’s and not their father’s genes, but many other examples are found across the animal kingdom. Extensive theoretical work has argued that such non-Mendelian reproduction could evolve as a result of conflict between the sexes, but empirical tests are sorely needed to test whether the plausible is actual.

To date, most empirical work exploring this conflict hypothesis has focused on mammals and other model organisms. Yet one of the most extreme and widespread case of non-Mendelian inheritance has barely been explored: in species with paternal genome elimination (PGE), males develop from fertilised eggs, but the complete haploid set of chromosomes inherited from their fathers is eliminated at some stage of development (Normark, 2003; Burt and Trivers, 2006; Gardner and Ross, 2014; Blackmon et al., 2015). PGE has evolved independently at least seven times across arthropods and is estimated to be present in more than 10,000 species (de la Filia et al., 2015). Under PGE, both sexes are diploid, in contrast to classic haplodiploidy—but, as in the latter, males only transmit maternally-inherited chromosomes to their offspring. Therefore, transmission of genes through males is not random, but dependent on whether they are maternally-derived. This provides a transmission advantage to mothers through their sons, and the evolution of PGE has been frequently framed as the outcome of intragenomic conflict between parental genomes in males—by maternal alleles being able to manipulate spermatogenesis in males to enhance their own transmission (Bull, 1979; Herrick and Seger, 1999; Ross et al., 2010; Normark and Ross, 2014). However, even if parental conflict was responsible for the initial evolution of PGE, it is unclear if this conflict is ongoing in extant species.

In addition to biased allelic transmission, PGE can also affect the chromatin state of paternal chromosomes, which has been suggested to result in extreme parent-of-origin-dependent gene expression (genomic imprinting). In several species where elimination of paternal chromosomes is postponed until spermatogenesis (germline PGE)—such as mealybugs, coffee borer beetles, and the booklice *Łiposcelis*—paternal chromosomes are heavily condensed and compacted into a dense body at the periphery of the nucleus, reminiscent of the mammalian bar body (Borgia, 1980; Nur, 1990; Hodson et al., 2017). This process takes place during embryogenesis, and, as a result, males are thought to become functionally haploid, only expressing maternal alleles (Prantera and Bongiorni, 2012). Like other cases of parent-of-origin specific gene expression, such as genomic imprinting of single genes in mammals and flowering plants (Reik and Walter, 2001; Ferguson-Smith, 2011), this is thought to result from intragenomic conflict between maternally and paternally derived alleles: silencing of the paternal genome in males could prevent it from expressing anti-PGE adaptations to escape elimination and restoring fair Mendelian transmission or from reducing male fitness under sibling competition (Herrick and Seger, 1999; Ross et al., 2010, 2011). Although paternal genome silencing has evolved repeatedly in the context of PGE, the hypothesis that it has evolved as an outcome of genomic conflict remains to be tested.

In order to address this, we study paternal genome silencing in mealybugs (Hemiptera; Pseudococcidae), small hemipteran plant-feeding insects that reproduce through PGE. We present the results from a RNA-seq allele-specific expression analysis (ASE) (Wang and Clark, 2014) in male offspring of hybrid and intraspecific crosses allowing us to detect and quantify parent-of-origin effects on gene expression at a whole-genome scale. We focus on three key questions: first of all, is the paternal genome completely inert, or do some paternal genes escape silencing? Previous work suggesting the completely silenced state of the paternal genome relied on cytogenetic observations or expression of a small number of phenotypic traits or genetic markers (Brown and Nur, 1964; Brown and Wiegmann, 1969; Brown, 1972; Brun et al., 1995; Borsa and Kjellberg, 1996), and therefore lack the genome-wide resolution to address this question. Complete silencing of paternally-inherited chromosomes would indicate that there is very little scope of ongoing intragenomic conflict within mealybug males; however, any genes that escape silencing might be involved in conflict (Herrick and Seger, 1999; Ross et al., 2010, 2011). Second, is paternal genome silencing restricted to somatic tissue, or does it also occur in the germline? Paternal chromosomes are eliminated during spermatogenesis, so we might expect the testis to be a hotspot for intragenomic conflict and experience stronger selection for silencing of the paternal genome. We test this hypothesis by comparing patterns of parent-of-origin expression between male soma and testis. Finally, do paternally-expressed genes evolve faster than genes with complete maternal expression? Evolutionary conflict can lead to arms races in which each party evolves rapidly in response to the harm inflicted by the other. As a result, genes involved in this conflict might evolve rapidly (Geist et al., 2019). We therefore test if genes that escape paternal genome silencing, particularly those in the testis, show an elevated rate of molecular evolution.

## Results

In order to determine whether paternal genomes retain transcriptional activity or are completely silenced in male mealybugs, we estimated patterns of allele-specific expression (ASE) in 1) somatic and germline tissues of adult hybrid males (CF males) originated in crosses between citrus mealybug (*Planococcus citri*) females and males from the closely related vine mealybug (*P. ficus*), and 2) whole adult *P. citri* males produced in reciprocal intraspecific crosses between three pairs of isofemale lines (WYE3-2 x CP1-2, WYE3-2 x BGOX-6 and CP1-2 x BGOX-6). A detailed overview of experimental methods and analysis strategy is given in Materials and Methods.

### Allele-specific expression in hybrid males

We obtained transcriptomes from three biological replicates of soma and testes dissected from pools of CF hybrid males (113-158M reads per sample) (Fig. 1A) and aligned them to a custom pseudogenome constructed from the PCITRI.V0 reference genome (https://ensembl.mealybug.org/) and a panel of 4,670,197 interspecific SNPs between the parental *P. citri* and *P. ficus* genomes (Materials and Methods). We found and validated 159,059 and 221,600 SNPs from the interspecific panel in all three CF soma and testis transcriptomes, respectively, at a read depth ☒ 30 (Fig. S1). At each SNP, we estimated the proportion of reads that originated from the maternal genome (p_m_) and found that the vast majority of reads are of maternal origin in both soma (0.88±0.13) and testes (0.97±0.06) (Fig. 1B, Table S1), as expected under somatic heterochromatinisation of paternal chromosomes.

**Fig 1.**
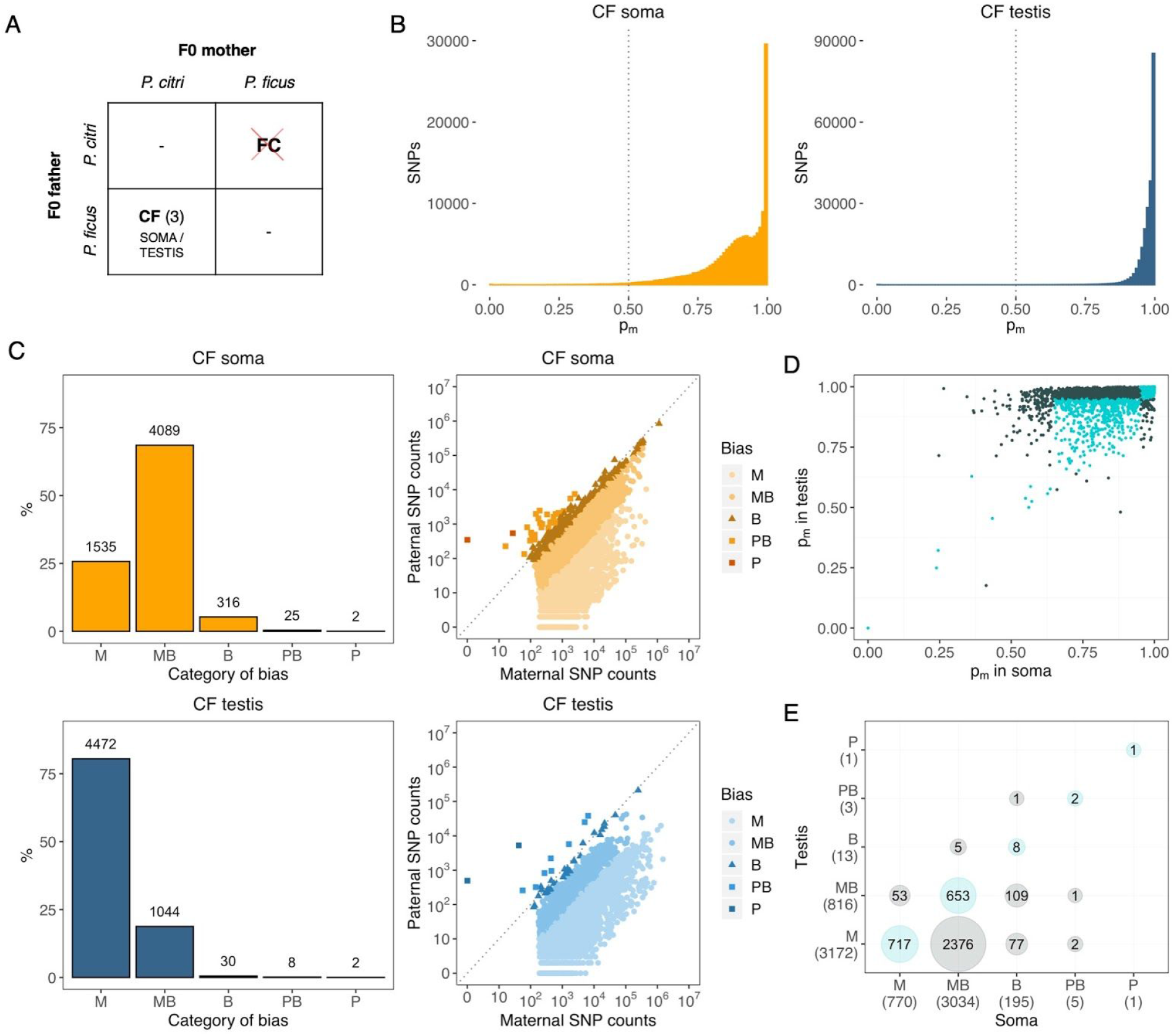
Quantification of allele-specific expression (ASE) in soma and testis of hybrid F1 mealybug males. (**A**) Cross scheme between *P.citri* and *P. ficus*. Only CF crosses prcduced viable adult male offspring (number of replicates in brackets) Testes from CF males were dissected and sequenced separately from the soma (**B**) Histogram of expression biases to maternal genome, p_m_, at SNP level in soma (orange, left) and testis (blue, right) of F1 mealybug males (pooled maternal and paternal counts between replicates). The dotted line indicates complete biparental expression. (**C**) Counts of genes with allele-specific information according to category cf ASE expression (from completely maternal, M, to completely paternal, P) (left panels) and scartterplots of combined paternal and maternal counts (across exonic SNPs and replicates) for genes expressed in soma and testis (right panels). (**D**) Scatterplot of p_m_ in soma and testis for all genes with allele-specific information expressed in both tissues (genes belonging to the same ASE category in both are shown in turquoise). (**E**) Cross table of overlapping gene counts by ASE category in soma and testis.

To estimate ASE at gene level, we assigned 93,039 SNPs in soma (59% of total) and 101,333 in testis (46%) to coding regions in the reference genome. Out of 18,667 genes with detectable expression levels (TPM>1) in soma and 15,286 in testis, we were able to confidently estimate ASE in 5,967 and 5,556 genes, respectively. These genes are those covered by at least two SNPs in all three tissue replicates (or a single SNP with average read depth > 100), and showing homogeneity in expression bias across tissue replicates (Materials and Methods). To generate gene-level ASE estimates for each of these genes, we pooled maternal and paternal read counts across all exonic SNP (on average, 14.7 SNPs per gene) in each individual replicate and estimated genic p_m_ as the fraction of reads originated from the maternal genome. Since soma and testis replicates show high consistency in genic p_m_ estimations (average p_m_ difference between replicates of |0.026| and |0.017|, respectively), we obtained a single p_m_ estimate for each gene in soma and testis by pooling maternal and paternal counts from all three replicates.

Concordantly with patterns observed at SNP level, gene expression is globally biased towards the maternal genome in both soma and testis (Fig. 1C). In soma, only 5.2% of genes (316) exhibit substantial biparental (B) expression, defined (after Wang and Clark, 2014) as p_m_=0.35-0.65 and/or Bonferroni-corrected exact binomial test vs p_m_=0.5 not rejected. The majority of somatic genes (68.5%, 4,089 genes) are predominantly expressed from the maternal *P. citri* genome (maternally-biased, MB, with 0.65<p_m_<0.95) and a further 25.7% of genes (1,532) are completely maternal (M, p_m_≥0.95). Additionally, we found 25 somatic genes that are predominantly expressed from the paternal *P. ficus* genome (PB, 0.05<p_m_<0.35) and 2 completely paternally-expressed genes (P, p_m_≤0.05).

In testis, ASE patterns are significantly different to those found in soma (G-test of independence, G=3,700.5, df=4, P<0.001) and even more shifted towards the maternal genome, with >99% genes showing a predominance of maternal expression: 80.5% (4,472) are completely M, and 18.8% (1,044) MB. Only 0.5% of testis-expressed genes (30) can be classified as B, while we found 8 and 2 genes to be, respectively, PB and P.

The observed differences between tissues are indicative of different patterns of global allele-specific expression in soma and germline. We also analysed differences in ASE for the 4,005 overlapping genes which were expressed in both soma and testis and found only moderate correlation (Spearman’s ρ=0.50) between p_m_ estimates in both tissues (Fig. 1D). Only 34.5% belong to the same ASE category in both tissues (Fig. 1E), which is indicative of changes in imprinting status between somatic and germline tissues, with paternal chromosomes contributing to transcription at a greater extent in the soma.

### Parent-of-origin expression in intraspecific males

In order to confirm that the global bias to the maternal genome observed in CF males is due to parent-of-origin effects (which cannot be determined without the reciprocal cross), and to rule out that the extreme genome-wide imprinted expression pattern is a product of hybridisation, we also estimated ASE in pure *P. citri* males produced in reciprocal crosses between three isofemale lines (Fig. 2A).

**Fig 2.**
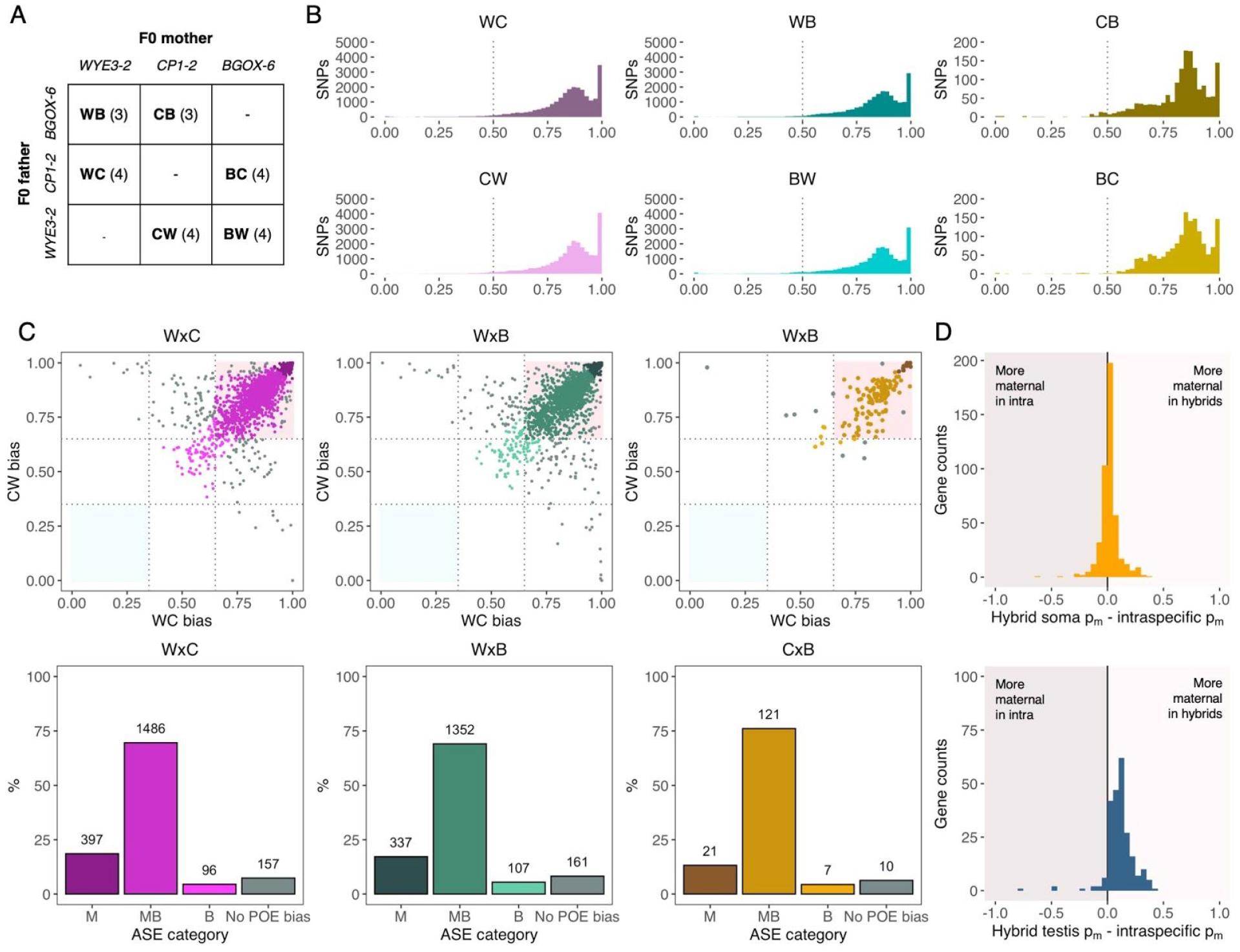
Quantification of parent-of-origin ASE in F1 *P. citri* males from pairs of reciprocal crosses between three isofemale lines. (**A**) Cross scheme between the three lines (number of replicates in brackets) (**B**) Histogram of expression biases to maternal genome, p_m_, at SNP level in all six genotypes (pooled maternal and paternal counts between replicates). The dotted line indicates complete biparental expression. (**C**) Gene p_m_ scatterplots in both reciprocal genotypes from each parental cross pairs (lop panels). Maternally-biased genes are expected in the top right comer (pink), while paternally-biased genes are expected in the bottom left panel (light blue). Genes with discordant p_m_ in reciprocal genotypes (not showing a true parent-of-origin ASE pattern) are shown in gray. Gene counts divided by ASE category are shown in the bottom panels. (**D**) Histogram of p_m_ differences between soma-only genes (top panel)Ztestis-only genes in CF hybrids and intraspecific males (p_m_ averaged across all three parental cross pairs).

We found 24,660 high-confidence SNPs shared with a read depth ☒ 20 in the transcriptomes of 4 WC and 4 CW replicates originated in reciprocal crosses between WYE3-2 and CP1-2 (WxC), 20,903 SNPs in 3 WB and 4 BW replicates from WYE3-2 x BGOX-6 (WxB) crosses and only 1,592 SNPs in 3 CB and 4 BC replicates from CP1-2 x BGOX-6 (CxB) crosses (Fig. S2) (107-218M reads per sample). As in hybrid crosses, p_m_ distribution across SNPs shows a strong bias towards the maternal genome in all genotypes (Fig. 2B), with averages ranging between 0.82-0.85 (0.14-0.15) (Table S2). We assigned 16,748 exonic SNPs (68% of total) to 2,136 genes in WxC replicates, 14,775 (71%) to 1,957 genes in WxB replicates and 1,054 (66%) to 159 genes in CxB replicates (6.8 SNPs per gene in WxC and WxB crosses and 5.7 in CxB crosses). Assignment of genes to ASE categories for each of the six intraspecific genotypes individually revealed similar patterns to those observed in CF soma, with a predominance of MB and M genes (Fig. S3).

Actual parent-of-origin ASE patterns could be estimated by crossing p_m_ estimates between each pair of reciprocal genotypes (Fig. 2C). Average p_m_ differences between reciprocal crosses are small in all three pairs (|0.055|-|0.065|), with 85-87% of genes belonging to the same ASE category in both cross directions. Incorporating reciprocal information allowed us to determine that only 6-8% of genes show allelic biases that are not consistent in both cross directions and therefore do not represent true parent-of-origin effects (“no POE bias”). These genes include all those that had been classified as predominantly paternally-expressed (PB or P) in individual genotypes, suggesting that the putatively PB and P genes found in hybrid soma and testis do not represent true parent-of-origin expression and are most likely under line-specific effects. Among the genes with consistent parent-of-origin effects in intraspecific crosses, 69-76% exhibit MB expression and a further 13-19% are completely M, with only 4-5% showing biparental expression. These patterns are consistent across all three intraspecific crosses (G-test of independence, df=6, P=0.26). p_m_ estimations show high consistency (Spearman’s ρ=0.83) between intraspecific and hybrid males for 461 soma-limited genes (i.e. present in intraspecific and CF soma datasets but not in CF testis), but only moderate for 231 testis-limited genes (ρ=0.35), which showed a higher bias to the maternal genome in hybrids (Fig. 2D). Overall, the data in both hybrids and intraspecific crosses reveal a consistent parent-of-origin, tissue-specific global expression bias to the maternal genome consistent with cytogenetic observations of paternal genome heterochromatinization (see below), albeit with partial transcriptomic activity of paternal chromosomes.

### Cytological evidence of paternal genome silencing in the germline

We complemented our transcriptomic analysis of allele-specific gene expression in hybrid testis with a cytological time-series of paternal genome condensation dynamics in *P. citri* males in successive larval stages (Fig. 3). In mealybugs, spermatogenesis begins at 2nd larval instar (Nur, 1962; Bongiorni et al., 2004; Bain, 2019). The dense nuclear bodies which result from heterochromatinisation of paternal chromosomes can be first observed in the rapidly dividing somatic cells of the developing accessory glands (AG) in the 2nd larval instar (Fig. 3B-D), coinciding with early spermatogenesis. During 3rd instar (late spermatogenesis), all cells of the fully developed AG clearly show heterochromatinisation of paternal chromosomes. Interestingly, however, during 4th instar paternal chromosome heterochomatinisation can only be observed in a fraction of the cells of the AG, and the signal is completely lost in the adult testis. In addition to germline tissues, we also targeted Malpighian tubules, which have been shown to lack paternal genome heterochomatisation in larval male instars (Nur, 1967). The dynamic patterns of paternal chromosome silencing and decondensing in testis are in stark contrast to Malphigian tubules, which consistently lack or show very limited paternal chromosome heterochromatinisation across all male stages (Fig. 3E). Since our testis allele-specific expression patterns are consistent with strong paternal chromosome heterochromatisation, we speculate that there is very limited transcription taking place in the adult testis (as spermatogenesis has already finished in adults) and our RNA-sequencing captured residual gene expression from earlier stages.

**Fig 3.**
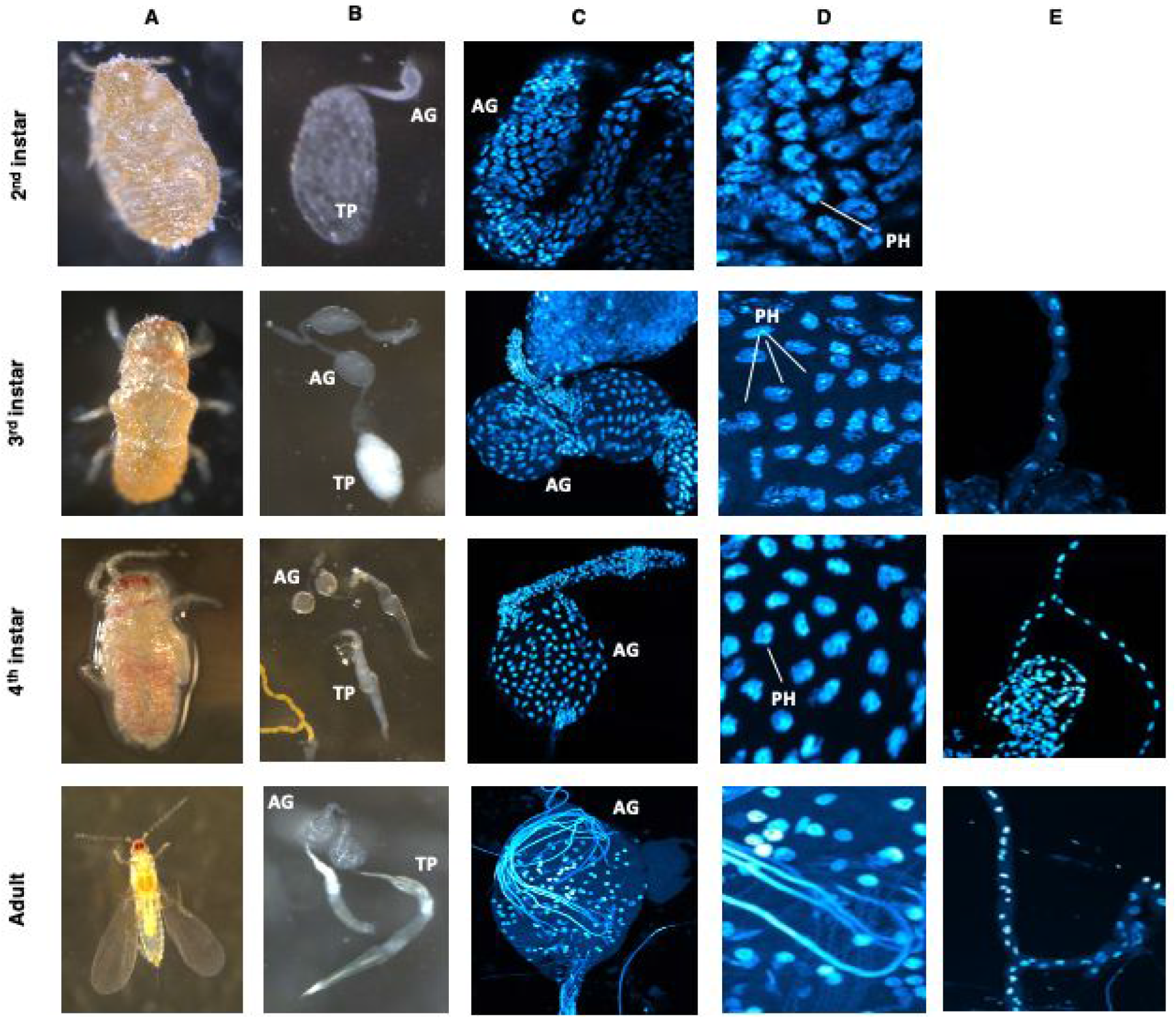
Cytogenetic analysis of paternal genome heterochromatization in cells of the accessory glands and testis proper end Malpighian tubes at 4 stages of development: 2nd instar, 3rd instar, 4th instar and adult. AG, accessory gland; TP, testes proper; PH, paternal genome heterochromatization. (**A**) images taken on a dissection microscope of the individual from which the testes and Malpighian tubes were dissected. (**B**) Images taken on a dissection microscope of the dissected testes. (**C**) Confocal image of the accessory gland stained with DAPI (x63). (**D**) Enlarged section from image (**C**) showing PH status. (**E**) Confocal image of the Malpighian tubes stained with DAPi (x63)

### Functional investigation of genes without maternal expression

We then examined the functional profile of the minority of genes exhibiting biparental or predominantly paternal expression in hybrids and intraspecific males identified in our ASE analysis. In hybrid soma, where our detection power is highest thanks to more genes with allele-specific information, 172 B, PB and P genes have at least a GO term assigned, against a background population of 3,193 genes with ASE information and GO assignment. We found four GO terms significantly enriched in biparental and paternally-biased genes (FDR<0.05): translation, oxidation-reduction process and ribosome/structural component of ribosome (Table S3). We did not find significant enrichment for any GO terms in hybrid testis (172 B, PB and P genes GO-annotated versus a population of 1,535) or intraspecific males (77/1,231).

We further examined putative functions of biparentally-expressed genes using BLAST homology. We found genes with mitochondrial functions to be common among B and PB genes in hybrid soma (100/292 annotated genes, Dataset S1) and intraspecific (29/108, Dataset S2). These include genes that code for structural components of mitochondrial ribosomes: for example, in hybrid soma, 6 small subunit proteins, mRpS2, mRpS12, mRpS15, mRpS23, mRpS29 and mRpS30, and 19 large subunit proteins: mRpL1, mRpL3, mRpL4, mRpL9, mRpL10, mRpL12, mRpL18, mRpL19, mRpL20, mRpL22, mRpL24, mRpL27, mRpL28, mRpL32, mRpL37,mRpL38, mRpL39, mRpL43 and mRpL52) and members of the mitochondrial respiratory chain, such as homologs to *D. melanogaster* ND-18, ND-20, ND-23, ND-30, ND-39, ND-49, ND-51, ND-B14 and ND-PDSW (NADH:ubiquinone oxidoreductase complex I), ATPsynF and ATPsynO (mitochondrial ATP synthase complex V) or Cyt-c-p (cytochrome c). We also found several B genes related to glucose metabolism (6-phosphofructokinase, inositol-3-phosphate synthase, ribulose-phosphate-3-epimerase) and fatty acid metabolism (fatty acid synthases, short-chain synthetase and ligase, long-chain ligase and dehydrogenase) in both hybrid soma and intraspecific males. In contrast, these genes with mitochondrial functions identified in hybrid soma and intraspecific males were found to be M or MB in hybrid testes, where only 25 B genes could be annotated (Dataset S3). Among these genes, we did not identify any candidates involved in reproductive functions.

Since many of the above B genes identified hybrid soma and intraspecific males are constitutive, we speculated that biparentally expressed genes would tend to exhibit higher expression levels than completely maternal genes. To assess the effect of bias to the maternal genome in gene expression levels, we fitted a linear model with log-transformed TPM counts as the response variable and p_m_ and the quadratic term p_m_^2 as fixed effects. In order to obtain independent estimates of gene expression level and bias to maternal genome, we used gene TPM counts estimated from three whole adult *P. citri* transcriptomes (see Materials and Methods) and p_m_ estimates obtained from intraspecific F1 males. We found both a significant linear and quadratic relationship between p_m_ and gene expression, albeit with a very low effect size (R^2^ adjusted = 0.026, F_2,2376_ = 33.42, P < 0.001, Fig. S4).

Also, in addition to measuring parent-of-origin expression, our RNA-sequencing design allowed us to examine more straightforward sex-specificity of gene expression. We evaluated whether biparental genes are more male-specific than genes with higher degree of maternal expression using a specificity metric (SPM)—a proportion of expression ranging from 0 (male-specific) to 1 (female-specific)—calculated from transcriptomes of whole *P. citri* males and females (Materials and Methods). We fitted a GLM to explore the relationship between SPM and parent-of-origin expression in intraspecific F1 males and found a extreme distribution of sex-biased genes in the *P. citri* transcriptome (Fig. S5), which might be driven by the extreme sexual dimorphism in mealybugs. Both p_m_ (Z = 12.26, p < 0.001) and p_m_^2 (Z = −12.89, p < 0.001) were significant, indicating that both biparental and completely maternal genes tend to be more male-specific than genes showing an intermediate degree of bias to the maternal genome. (Fig. 4A)

**Figure 4:**
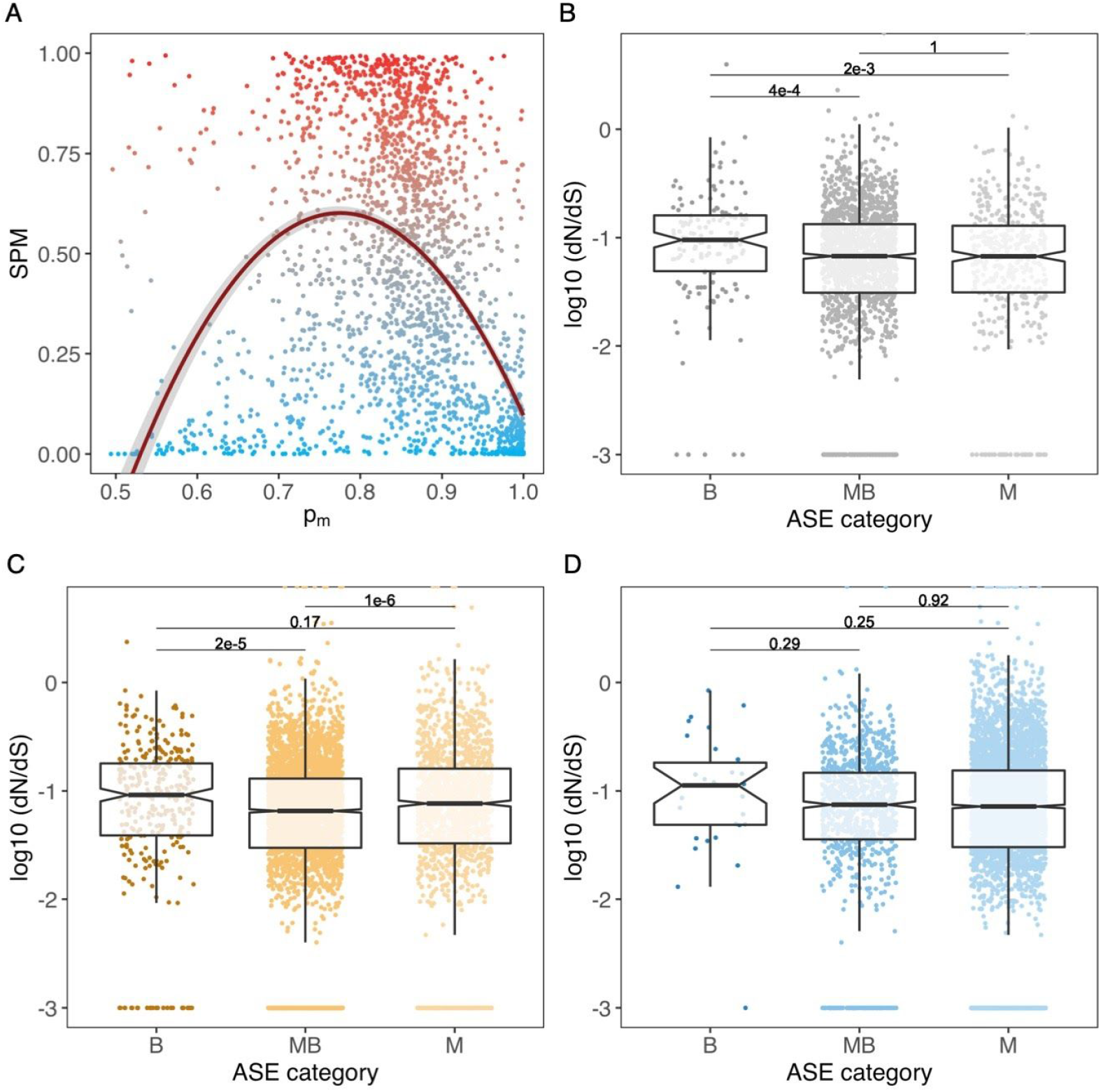
Sex-specifiC expression and evolutionary rates of genes with consistent pareet-of-origin information in intraspecific males. (**A**) Scatterplot of sex-specificity and bias to the maternal genome p_m_ for all 2,197 genes with consistent POE patterns. Transcriptomes of *P. citri* males and females from the WYE3-2 line were used to estimate sex-specificity (SPM), (blue dots represent male-biased genes, grey, non-biased genes; red, female-biased genes). The regression curve predicted by a GLM exploring the relationship between sex-specificity and bias to the maternal genome (SPM ~ p_m_ + p_m_^2) is shown in dark red. (**B**) Eox plot showing differences in evolutionary rates (as log-transformed dN/dS ratios) between biparental, materially-biased and maternal-only expression in intraspecific crosses. The p-values cf post-hoc Nemenyi tests in pairwise comparisons between ASE categories are shown. To plot genes with dN/dS = 0, a 0.001 constant was added to all dN/dS values (**C**) dN/dS ratios between biparental, maternally-biased and maternal-cnly genes in hybrid soma. (**D**) dN/dS ratios between biparental, maternally-biased and maternal-only genes in hybrid testes.

### Rates of molecular evolution between biparental and maternal genes

Our interspecific crossing design presented an opportunity to examine long-term evolutionary rates of genes with different parent-of-origin expression patterns. We aligned *P. ficus* sequences to the *P. citri* reference genome to obtain variants and annotated the effects (i.e. synonymous or non-synonymous) with a pipeline described Materials and Methods. Due to the slight inconsistencies in ASE bias between reciprocal crosses for some genes, we only considered biparental genes with consistent parent-of-origin effects in intraspecific F1 males. In these, we found a significant effect of expression type (B, MB or M) on rates of molecular evolution (dN/dS) using a non-parametric Kruskal-Wallis test (χ^2^_2_= 14.74, p = 0.0006). Using a post-hoc Nemenyi test, this difference is driven by elevated dN/dS in biparentally expressed genes, which evolve faster than maternally-biased (p = 0.0004) or maternal-only (p=0.002) genes. The two classes of maternally skewed genes do not evolve differently from each other (p = 1.000) (Fig. 4B).

We also repeated these analyses with parent-of-origin expression in soma and testes in CF males, to examine if these differences in evolution rates were also found in hybrids and specifically in germline tissues. While we also found a significant effect of expression class in dN/dS ratios in hybrid soma (χ2_2_= 40.23, p < 0.001), biparental and maternal-only genes both evolve faster than maternally-biased genes but not differently from each other (Fig. 4C). In hybrid testes, however, biparental, maternally-biased and maternal only genes do not show significant differences in their evolution rates (χ2_2_= 2.62, p = 0.270) (Fig. 4D).

## Discussion

Mealybugs have a unique reproductive strategy, where males inherit a haploid genome from both parents, but the paternally-inherited chromosomes are eliminated during spermatogenesis and excluded from viable sperm. This type of reproduction is thought to both originate from and lead to strong intragenomic conflict between maternally- and paternally-inherited alleles. Transcriptional repression of the paternally-inherited genome in males has been suggested to alleviate this conflict, yet had not been studied on a genome-wide scale.

In this study, we show that global patterns of gene expression do indeed show a bias towards expression of maternal alleles: using an allele-specific expression analysis, we estimated allele-specific expression profiles for >5,500 genes in somatic and reproductive tissues of hybrid mealybug males originated from crosses between *P.citri* females and *P. ficus* males. We found a strong predominance of maternal gene expression in both soma and testis, with up to ~95% and ~99% genes, respectively, showing evidence of bias towards the maternal genome. We confirmed these results in subsequent analysis of male offspring from intraspecific *P. citri* males, where >2,100 genes reflected the strong bias to the maternal genome observed in hybrids. We show that parent-of-origin expression profiles are extremely consistent, not only between biological replicates but also across genotypes. Thus, we conclusively show a strong parent-of-origin effect in gene expression in mealybugs whereby maternal chromosomes are the main contributors to transcriptomic activity in males.

However, despite this global maternal bias in gene expression, we find that the paternal genome is far from inert and that in fact paternally-inherited alleles are expressed to some extent for the majority of genes. Paternally-inherited copies escape silencing to some degree across approximately ~75-80% of genes in hybrid soma and whole intraspecific males. This percentage is lower is male testes, where only ~20% of genes are expressed with some contribution of the paternal allele. This difference between degree of paternal genome silencing between soma and testis suggests that paternal alleles are expressed in a tissue-specific manner. For example, many genes with exclusive maternal allelic expression in the testis show incomplete silencing in the soma. In mealybug males, the paternally-inherited chromosome set becomes highly condensed during early embryogenesis and is enriched for repressive histone marks, such as H3K9Me3 (Bongiorni et al., 2007). It seems unlikely that a large number of paternally-inherited alleles can be transcribed from such heavily condensed chromosomes, especially as there is direct evidence of RNA synthesis inhibition from the heterochromatic set (Berlowitz, 1965). Therefore, expression of paternal alleles is most likely due to tissue or cell-specific loss of heterochromatinisation. This was first suggested by Nur (1966, 1967, 1970) based on cytological observations showing that embryonic condensation of paternally-inherited chromosomes is later reversed in cells of a small number of male tissue types, such as the Malpighian tubules. We confirm these early observations in this study, and also show that the paternal genome condensation state can vary within the same tissue/cell type across development. We therefore hypothesize that genes showing an intermediate degree of bias towards the maternal genome are expressed in cells with and without heterochromatinisation of paternally-inherited chromosomes, while biparentally-expressed genes are probably specific to tissues without heterochromatinisation, or sufficiently enriched in those tissues to dominate the allele-specific expression signal. Of course, it is also possible that some genes are still expressed from heterochromatic chromosomes. The best known example of whole chromosome silencing, X-inactivation in mammals, is also achieved via heterochromatinisation (Lyon, 1961; Wutz, 2011). However, a small proportion of genes (1-3% in mouse, 15% in human) on inactive X chromosomes in females escape inactivation to some degree (Carrel and Willard, 2005; Filippova et al., 2005; Heard and Disteche, 2006; Yang et al., 2010; Crowley et al., 2015). This might also occur in mealybugs, especially for those constitutive genes (such as mitochondrial ribosomal proteins) which we found to be biparentally expressed in all tissues.

In three of the 7 independent origins of PGE described so far in arthropods—including mealybugs—whole-genome meiotic drive of maternally-inherited chromosomes has led to the subsequent evolution of silencing of the paternally-inherited set through heterochromatisation (Burt and Trivers, 2006; Gardner and Ross, 2014; Hodson et al., 2017). This recurring evolution of transcriptional suppression of the paternal genome has been interpreted as a response to the strong intragenomic conflict within males generated by PGE, as they only transmit their maternally-inherited genome to their offspring (Herrick and Seger, 1999; Ross et al., 2010). In mealybugs, the maintenance of paternal chromosome silencing depends directly on the maternally-inherited set (Nur, 1962; Chandra, 1963; Brown and Nur, 1964), which has been interpreted as support for this hypothesis. But how can we explain the observed patterns of reactivation of the paternal genome? One possibility is that it results from an ongoing evolutionary arms-race between maternally- and paternally-inherited alleles. To test this hypothesis, we focused on expression patterns in the testis, as this is where elimination of paternal chromosomes takes place. Therefore, reactivation of paternal alleles in the testis might represent a paternal response to fight their exclusion from sperm (Herrick and Seger, 1999; Ross et al., 2010). Depending on the current dynamics of the arms race, testes could either be a hotspot for paternal chromosome reactivation or a tissue in which the control exerted by the maternal genome over their paternal counterpart is most stringent. Our analysis showing stronger silencing of the paternal genome in testis compared to soma tentatively supports the latter scenario.

However, it must be noted that male meiosis and the subsequent degradation of spermatids containing paternal chromosomes mostly takes place prior to adulthood, during the second and third larval instars (Nur, 1962; Bongiorni et al., 2004; Bain, 2019). By targeting adult male testes for RNA-seq, we might have missed paternally-expressed genes acting during or just prior to male meiosis. We show cytological evidence of widespread heterochromatinisation of paternal-inherited chromosomes in testis of 3rd instar males, and also observed heterochromatinisation of paternal chromosomes in some testis cells in 2nd instar males—but, unfortunately, how ubiquitous it is during this stage is less clear, due to the fact that germline cells undergo rapid cell division during this stage. Therefore, future RNA-seq analyses of these developmental stages would be particularly informative.

The seemingly specific biparental expression of functionally important genes further suggests that the maternal genome may be firmly in control of the paternal genome, allowing its expression only for subsets of genes that are important for male function and do not pose a risk to the interests of the maternal genome (i.e. loci at which conflict is less likely to arise). Reactivation of the paternal genome might then be selected for to increase transcription of particular genes (as first suggested by Nur, 1966). In hybrid soma (where our power to quantify allele-specific expression is highest), biparentally-expressed genes are enriched in fundamental processes such as translation and oxidation-reduction. Many of these genes play important roles in the mitochondria, including mitochondrial ribosome proteins (MRPs) and others involved in the electron transfer chain, suggesting that reaching diploid expression levels could be important if mitochondrial function is limiting. MRPs are encoded by nuclear genes and imported into the mitochondria, where they are assembled into ribosomal subunits in conjunction with mitochondrially-encoded rRNAs (Richman et al., 2014; Tselykh et al., 2005; Rackham and Filipovska, 2014). Mitochondrial ribosomes are responsible for translation of mRNAs encoded by mitochondrial genes, including those involved in ATP production (O'Brien, 2002; Beckmann and Herrmann, 2015). Of course, a possible reason for the overrepresentation of MRPs genes might be their accelerated evolution rates (O'Brien, 2003; Desmond et al., 2011), which allows a more reliable estimation of allele-specific expression due to higher SNP density.

Interestingly, other biparental genes that we were able to identify in soma and whole males are involved in glucose and fatty acid metabolism. Mealybug males cease to feed at late 2nd instar (Franco et al., 2009), so upregulating transcription of gene pathways dedicated to energy production could be a mechanism to compensate for dietary restriction. One of the cell types that can undergo heterochromatisation reversal in *P. citri* are oenocytes (Nur, 1966), which are specialised in lipid storage and metabolism (Makki et al., 2014). Although we are constrained by limited homology to genes in better studied organisms such as *D. melanogaster* or *A. pisum*, our data suggest that paternal chromosomes may be recruited to boost transcription to diploid levels in a tissue-dependent manner for energy production and lipid metabolism, among other processes. In short, reversal of heterochromatinisation might be selected for to boost transcription in tissues with high metabolic demands.

Finally, in line with the idea that biparental expression is not the result of stochastic errors in gene silencing, we found that biparentally expressed genes are faster evolving (i.e. have higher dN/dS ratios) than genes with predominantly maternal expression in intraspecific males. However, this is not the case in germline tissues of hybrid males, where genes apparently evolve at the same rate—although our ability to detect differences in dN/dS rates in testes might be hindered by the limited number of biparentally-expressed genes identified in the testes (which is in itself suggestive of differences in strength of intragenomic conflict between the soma and germline). The implications of differences in evolutionary rate are difficult to parse with the current data. For instance, if paternal genes escaping silencing are indeed in an evolutionary arms race with the maternal silencing machinery, the faster evolution would be driven by positive selection of novel variants. However, faster evolutionary rates do not *per se* imply adaptation, as such patterns can easily arise from the opposite: relaxed selection, as has been argued for genes involved in reproduction (Dapper and Wade, 2016). It is worth noting that genes with maternally-biased and, especially, strictly maternal expression are necessarily expressed in the haploid state. As such, any deleterious variation, even that which would otherwise be recessive, is exposed to selection in males, resulting in strong purifying selection (Gerstein et al., 2011). From this perspective, the apparent elevation of dN/dS of biparentally expressed genes in intraspecific males might be more correctly viewed as a decrease in dN/dS of maternally-biased genes, thanks to an increased role of purifying selection. It will require more careful scrutiny of within-species variation to obtain estimates of adaptive evolution across parent-of-origin-bias categories and disentangle these alternative explanations.

## Conclusions

In this study, we show genome-wide parent-of-origin expression in an insect whereby maternally-inherited chromosomes are the main contributors to transcriptomic activity in males. In recent years, a number of studies have considered genomic imprinting beyond mammals and flowering plants (de la Casa-Esperón, 2012; MacDonald, 2012). We now have several allele-specific studies of imprinted gene expression (or lack thereof) in insects (Coolon et al., 2012; Kocher et al., 2015; Wang et al., 2016; Galbraith et al., 2016; Marshall et al., 2020). Yet most of these show very limited evidence for genomic imprinting, so our understanding of this process in insects remains limited. So far, most studies have focused on eusocial Hymenoptera, specifically to test the kinship theory of genomic imprinting, (Queller, 2003; Patten et al., 2014). However, the interactions between relatives with asymmetric genetic relationships that underlie this theory are not exclusive to classic haplodiploidy and eusociality. PGE taxa share the same genetic relationships between relatives as those found in true haplodiploids, but unlike in classic haplodiploidy males are still diploid and carry a complete haploid copy of their father’s genome, which is not transmitted to the offspring. As a result, there might be conflict between maternally- and paternally-inherited alleles over transmission—and over male fitness in scenarios of sibling competition, as paternally-inherited alleles in males might favour the fitness of female relatives over their own (Ross et al., 2011).

From an intragenomic conflict perspective, the differential parent-of-origin-specific expression patterns between somatic and reproductive tissues of mealybug males found in this study suggest an coevolutionary scenario over maintenance of paternal silencing in which the maternal genome appears to have stringent control over the paternal set, especially in testis. However, there are other potential explanations for these patterns that cannot be excluded. Targeting earlier larval stages, as discussed earlier, would allow capturing spermatogenesis directly and framing paternal genome expression patterns in the context of their elimination. Finally, another exciting prospect for this analysis strategy is its application to other germline PGE taxa with and without paternal chromosome silencing, in order to gain a comparative understanding of the somatic manifestations of this bizarre genetic system and its evolutionary implications.

## Materials and Methods

### Experimental crosses

Hybrid crosses were conducted between individuals from wild-derived, highly inbred laboratory strains: WYE3-2 (*Planococcus citri*, derived from an English population, >25 generations of sib-mating) and PF1-1 (*Planococcus ficus*, derived from an Israeli population, >10 generations of sib-mating). Due to extreme male specific mortality in hybrid offspring of crosses between *P. ficus* females and *P. citri* males, we were unable to raise viable adult male hybrids from these crosses. Therefore, only hybrid males from *P. citri* mothers and *P. ficus* fathers (CF crosses) could be sequenced. Intraspecific crosses were produced in all possible combinations between WYE3-2 and two additional isofemale *P. citri* lines, CP1-2 (derived from Israel) and BGOX-6 (derived from England) (>40 generations of sib-mating). All six reciprocal genotypes (WC, CW, WB, BW, CB, BC) yielded viable adult males.

Mealybugs were reared on sprouted potatoes placed on tissue paper in sealed plastic stock bottles and kept at 25°C and a 16h-light/8h-dark photoperiod without humidity control. Males and females used in experimental crosses were isolated before sexual maturity and isolated until adulthood. Hybrid crosses were set by placing pools (10-20 individuals) of brothers from the paternal species and sisters from the maternal species in 6cm-diameter glass Petri dishes. To encourage mating, a filter paper impregnated with 10 ng of synthetic sex pheromone from the paternal species (Bierl-Leonhardt et al., 1981; Hinkens et al., 2001) was placed in the Petri dish. After all males in the Petri dishes died, females were transferred to rearing bottles in groups to lay eggs. Hybrid F1 offspring were reared until becoming sexually differentiated (third instar). At that stage, viable CF males were transferred to glass vials sealed with cotton wool to reach adulthood. For intraspecific crosses, individual mating pairs were set in glass vials containing a single potato sprout. Successful matings were evidenced by females laying eggs 2-5 days postmating, which were transferred to individual bottles. Males were reared and isolated in glass vials until reaching adulthood.

### DNA and RNA extraction and sequencing

To sequence the genomes of the parental lines used in CF crosses, genomic DNA was extracted from 5-10 adult WYE 3-2 (*P. citri*) and PF1-1 (*P. ficus*) virgin females. Sample lysis, proteinase K digestion and RNA removal were performed using a DNeasy Blood & Tissue kit (Qiagen, The Netherlands) and isolation of gDNA was carried out with a Wizard Genomic DNA Purification Kit (Promega, USA) according to manufacturer’s instructions. A single TruSeq library (350 bp insert size) for *P. citri* and two TruSeq libraries (350 bp and 550bp insert sizes) for *P. ficus* were generated and sequenced on a HiSeq 2500 instrument by Edinburgh Genomics (The University of Edinburgh, UK). For intraspecific crosses, genomic DNA was extracted using a custom protocol from 6-8 virgin females from WYE3-2, CP1-2 and BGOX-6 sisters to the females used in the crosses and TruSeq DNA PCR free gel library (350 bp insert size) was sequenced on a HiSeq X instrument (150-bp read pairs).

For RNA-seq of CF males, we extracted RNA from somatic tissues and testes of >70 adult F1 male offspring from three pools of *P. citri* sisters mated to *P. ficus* males. Males were dissected in RNAlater (Thermo Fisher Scientific, USA) and soma and testes were immediately transferred to ice-cold TRIzol (Invitrogen) and stored at −80°C. RNA was extracted using isopropanol and chloroform (2.5:1) and linear acrylamide as a carrier. After extraction, residual gDNA digestion was performed using DNAse I (Thermo Fisher Scientific, USA) and RNA samples were purified with RNA Clean & Concentrator™−5 (Zymo Research, USA). Due to low yields of RNA for soma and testis CF samples, cDNA amplification was performed using the Ovation^®^ RNA-Seq System V2 (NuGen, USA). Two independent cDNA amplifications from each soma and tissue sample were performed separately to be sequenced as technical replicates. cDNA samples were purified using MinElute^®^ Reaction Cleanup Kit (QIAGEN, The Netherlands) in TE buffer. In total, 12 Illumina TruSeq Nano libraries (350 bp insert size) were generated and sequenced on two lanes on a Illumina HiSeq 4000 instrument (75 bp paired-end reads) by Edinburgh Genomics.

We also generated three Illumina TruSeq stranded mRNA-seq libraries from three pools of whole adult *P. citri* males and females from the maternal line used in the hybrid crosses (WYE3-2). The first male and female transcriptomes were sequenced on the Illumina HiSeq 4000 instrument (75 bp paired-end reads) and the remaining two on the Illumina NovaSeq S2 instrument (50 bp paired-end reads).

For intraspecific crosses, RNA was extracted from pools of 20-50 flash frozen full F1 brothers descending from a single cross using the PureLink RNA purification kit with DNase I digestion (Thermo Fisher Scientific, USA). TruSeq stranded mRNA-seq libraries (350 bp insert size) were generated from 23 samples (4 WC, 4 CW, 3 WB, 4 BW, 4 CB and 4 BC) and sequenced on a single lane of NovaSeq S2 instrument (50 bp paired-end reads) by Edinburgh Genomics.

DNA and RNA samples were quantified using Qubit BR Assay Kit (Thermo Fisher Scientific, USA) and their integrity was assessed via 1% agarose gel electrophoresis or Bioanalyzer RNA 6000 Nano kit (Agilent). Library quantification, normalisation and quality control were performed by Edinburgh Genomics (The University of Edinburgh, UK).

### SNP calling

To call discriminant SNPs between *P. citri* and *P. ficus*, we mapped 35.8M *P. citri* read pairs (16x coverage) and 123.3M *P. ficus* read pairs (40x coverage) to our reference *P. citri* genome (PCITRI.V0) with bwa 0.7.15-r1140 (BWA-MEM algorithm) (Li, 2013). We called a raw set of variants using FreeBayes v1.2.0 (Garrison and Marth, 2012) with the following settings: ---haplotype-length 0 --standard-filters ---min-alternate-fraction 0.05 -p 2 ---pooled-discrete --pooled-continuous. We used vcffilter (https://github.com/vcflib/vcflib#vcffilter) to filter the resulting VCF file (‘DP > 10 & SAF > 2 & SAR > 2 & RPR > 1 & RPL > 1’) and discard all non-SNP variants. We obtained an initial set of 5,288,538 discriminant SNPs between both genomes using the SelectVariants walker in GATK v3.8 (McKenna et al. 2010) to keep only variants that were monomorphic for an alternative allele in *P. ficus* and for the reference allele in *P. citri*. To keep only high-confidence sites, we then filtered out sites with AO < 20 and AO / DP < 0.99 to yield a final set of 4,670,197 between-species discriminant SNPs.

The same strategy was followed to call discriminant SNPs between the three pairs of isofemale *P. citri* lines used in intraspecific crosses. We mapped 183.7M read pairs for WYE3-2, 163.8M for CP1-2 and 325.0M for BGOX-6 to the reference PCITRI.V0 (81x, 77x and 135x coverages, respectively), called variants with freebayes and filtered the resulting VCF file with vcffilter as above. Then, we generated six separate VCF files (two for each pair of lines) in which the first line was monomorphic for the reference allele and the second was monomorphic for an alternate allele with DP > 10 in both. In total, we obtained 806,554 discriminant SNPs between WYE3-2 and CP1-2, 837,922 between WYE3-2 and BGOX-6 and only 57,154 between CP1-2 and BGOX-6.

### RNA-seq mapping and ASE estimation at discriminant SNPs

To estimate allele-specific expression patterns in male transcriptomes, we mapped RNA-seq reads to custom pseudogenomes for each hybrid and intraspecific cross generated by hard-masking SNP positions with different parental alleles.

Initial quality control and read trimming of raw sequencing data were performed with FastQC v0.11.5 and Trimmomatic v0.36 (Bolger et al., 2014) with default settings. After trimming, we obtain on average 48.7M (6.6 SD) trimmed RNA read pairs for each biological and technical replicate of hybrid soma and 51.1M (5.2 SD) for each biological and technical replicate of hybrid testes. In order to avoid mapping biases to the reference when estimating allele-specific expression, we built a custom pseudogenome using the FastaAlternateReferenceMaker walker in GATK v3.8 (McKenna et al., 2010) to hard-mask 4,670,197 SNPs between *P. citri* and *P. ficus* in the PCITRI.V0 reference. We separately mapped reads from each lane and technical replicate for all three soma and testis biological replicates against the pseudogenome using STAR v2.5.2b (Dobin et al., 2013) in the two-pass mode, marked duplicates with Picard v2.17 (http://broadinstitute.github.io/picard) and quantified expression levels using RSEM v1.3.0 (Li and Dewey, 2011). Due to high consistency across technical replicates for all samples (Fig. S6), we decided to merge BAM files from both technical replicates to estimate ASE estimation for each biological replicate (113.1-157.8M reads per sample in each merged BAM file).

We used the ASEReadCounter walker in GATK v3.7 to retrieve maternal (*P. citri*) and paternal (*P. ficus*) allele counts at discriminant SNPs with the following settings: -U ALLOW_N_CIGAR_READS-minDepth 30 --minBaseQuality 20 --minMappingQuality 255. Then, we applied a series of filters to remove low-confidence SNP positions from the ASE dataset (Fig. S1). First, we only kept SNPs present in all three tissue replicates. Second, we removed SNPs where the sum of uniquely mapped reference and alternate allele counts was <90% of the total read depth at that position in at least one replicate. Third, we used the male *P. citri* transcriptomes to remove polymorphic sites within *P. citri* undetected during SNP filtering. To do so, we mapped RNA-seq reads from the three independently-sequenced pools of adult *P. citri* males to the pseudogenome, passed the merged BAM file through ASEReadCounter with default settings and removed SNPs in the hybrid ASE dataset where the proportion of reference alleles in pure *P. citri* males was <95%.

The same general procedure was followed to estimate ASE in intraspecific crosses, including generating pseudogenomes unique to each of the three pairs of *P. citri* lines. We aligned 85.1 (16.0 SD) million trimmed read pairs per F1 sample to the pseudogenomes and run ASEReadCounter as above, but allowing a lower minimum read depth (20) to account for the lower number of discriminant SNPs between parental lines. Then, we applied the first (SNPs present in at least 3 replicates) and second above filters to the intraspecific datasets to retain only high-confidence SNPs in the F1 transcriptomes (Fig. S2). After inspecting distributions of parental counts in F1 males, we discovered that one of the CB replicates exhibited a markedly distinct ASE distribution to the others, with ~98% SNPs showing only maternal (CP1-2) alleles (Fig. S7). This led us to conclude that the mother of this cross had been erroneously classified as a virgin and had already been fertilised by a CP1-2 male prior to isolation, so this sample was excluded from subsequent analyses.

### Gene ASE expression analysis

SNPs in hybrid and intraspecific transcriptomes were assigned to PCITRI.V0 genomic annotation features using bedtools v2.27.1 (Quinlan and Hall, 2010) and classified as exonic, intronic, intergenic or orphan (if located on an unannotated contig). For all predicted genes, we pooled maternal and paternal read counts across all exonic SNP in each sample and estimated gene ASE as fraction of reads expressed from the maternal genome (p_m_). Genes covered by a single SNP were removed from subsequent analysis unless the pooled read depth across groups of biological replicates (soma and testis in hybrids; genotypes in intraspecific males) was at least 100. We followed (Wang and Clark, 2014) to assign genes to ASE categories: for each gene, we conducted exact binomial tests (with Bonferroni correction) separately in all replicates against the null hypothesis of Mendelian expression (p_m_=0.5) and genes were considered biparental (B) when the null hypothesis was not rejected and/or 0.35?p_m_≤0.65. Significant genes were classified as exclusively maternal (M) if p_m_≥0.95, maternally-biased (MB) if 0.65>p_m_<0.95, paternally-biased (PB) if 0.05>p_m_<0.35 and exclusively paternal (P) if p_m_≤0.05.

Additionally, we performed a G-test of independence with Bonferroni correction for each gene to test whether ASE was homogenous across all groups of replicates (Wang et al., 2016). Genes that did not show significant heterogeneity across soma and testis samples (in hybrids) and genotype replicates (in intraspecific males) were immediately validated. Significantly heterogeneous genes were included in the final analysis only if all replicates agreed on ASE category and significance of exact binomial test. After removing the genes that failed to meet these criteria, we estimated a combined p_m_ at gene by pooling paternal and maternal SNP counts from all groups of replicates and performing a final exact binomial test for each gene with these pooled counts.

Finally, we discarded low-expressed genes (TPM<1 on average across replicates) and, for intraspecific crosses, we only kept genes present in both cross directions for each of the three combinations of parental lines. We incorporated reciprocal information in the intraspecific crosses by averaging p_m_ between each pair of reciprocal genotypes and classifying them into ASE categories using the p_m_ thresholds described above. Genes that differed in category of ASE between reciprocal genotypes and showed a reciprocal p_m_ difference higher than the average across validated genes for each pair were not assigned a final ASE category.

### Functional annotation of genes

We performed a gene ontology (GO) enrichment analysis using the Fisher exact test with Benjamini-Hochberg FDR to identify functional categories enriched in all annotated biparentally and paternally-biased genes in soma and testis of hybrid males and in whole intraspecific F1 males using GOATOOLS (Klopfenstein et al., 2018). For intraspecific crosses, we tested genes that had been classified as biparental in at least one of the crosses. To reduce bias of enrichment analysis (Timmons et al., 2015), the background gene population was restricted to the gene dataset with ASE information for each group.

Biparentally, paternally-biased and paternal-only genes were further investigated. In addition to a default blastp search against UniProtKB/Swiss-Prot with an e-value cutoff of 1e^-10^, we identified reciprocal orthologues in the proteomes of *Drosophila melanogaster* and *Acyrthosiphon pisum* using BLASTp v2.7.1+ (E-value≤1e^-25^) and a modified version of the rbbh.py script (https://github.com/DRL).

### Calculating sex-specificity of gene expression and rates of molecular divergence

In order to examine sex-specificity of gene expression, we calculated the *P. citri* specificity metric (SPM), a proportion of expression ranging from 0 to 1 in a focal set of tissues (Xiao et al., 2010; Kryuchkova-Mostacci and Robinson-Rechavi, 2017). In our case, the contrasts were male and female tissue, such that an SPM of 1 for females indicates 100% of gene expression in females, 0.5 is unbiased expression between the sexes, and 0 is 100% expression in males. This metric is less common than differential expression analyses, which conventionally uses a 1.5 fold difference in expression (which corresponds to a 0.7:0.3 SPM) to define sex-biased genes. However, this fold-difference analysis belies how truly distinct male and female gene-expression are, with most genes expressed uniquely in one sex or the other (Fig. S5).

To estimate rates of molecular evolution for the different classes of expressed alleles, we started with the set of interspecific SNPs between *P. citri* and *P. ficus* described above. Next, we created a custom database for *P. citri* in the program SnpEff (Cingolani et al., 2012)in order to annotate variants as synonymous or non-synonymous. To properly calculate the scaled divergence rate (dN/dS, formally defined as the number of non-synonymous changes per non-synonymous site divided by the number of synonymous changes per synonymous site), we required information about the number of synonymous and non-synonymous sites in the exons of each gene. We took the set of annotated genes (.gtf file) and the reference genome (.fasta) and used a series of custom R scripts to annotate the codon-degeneracy of each protein-coding nucleotide in the genome and summed these values per-gene. With these dN/dS values, we compared evolutionary rates across expression categories for which we had consistent results across replicates (maternal-only, maternally-biased and biparentally expressed genes) using a non-parametric Kruskal-Wallis test to examine whether or not there was an effect of expression type on evolutionary rate. Finding a significant result, we investigated which differences drove this pattern with a post-hoc Nemenyi test. These statistical tests were all completed in R.

### Microscopy

2nd, 3rd, 4th instar and adult *P. citri* males were dissected in a drop of phosphate buffered saline (PBS) on a microscope slide using a Leica dissecting microscope. Whole testes and Malpighian tubes were isolated from all individuals and images of whole testes were taken using a Leica S8 APO dissection microscope. Excess PBS was removed using a cotton bud and tissues were fixed directly on the slide in a drop of PFA:acetic acid (1% PFA, 45% acetic acid, 54% dH2O) for 5 minutes. Excess PFA was removed using a cotton bud and 25ul of Vectashield Antifade Mounting Medium with Dapi (Vector Laboratories) was added directly to the tissue and cover slips applied. Slides were sealed with nail polish and stored in the dark at 4°C. Fluorescent Z-stack images of all tissue samples were taken using a Leica TCS SPE-5 confocal microscope and then processed and merged using ImageJ (https://imagej.nih.gov/ij).

## Data availability

All RNA-seq and DNA-seq data are available at XXXXXXXXX and wet lab protocols and custom R scripts at github.com/agdelafilia/ASE_in_mealybugs. The PCITRI.V0 reference genome is publicly available at https://ensembl.mealybug.org/.

## Acknowledgements

The authors would like to thank Andrew Clark and Doris Bachtrog for initial discussions and advice on experimental design and Beatriz Vicoso, David Finnegan and Scott Roy for further discussion and comments. This work was supported by a Natural Environment Research Council Independent Research Fellowship (NE/K009516/1, to LR), a European Research Countil Starting Grant (PGErepro, to LR) and the Darwin Trust of Edinburgh (AGF).

