## Supplementary material for "Males that silence their father’s genes: genomic imprinting of a complete haploid genome": Supplememtary Figures and Tables

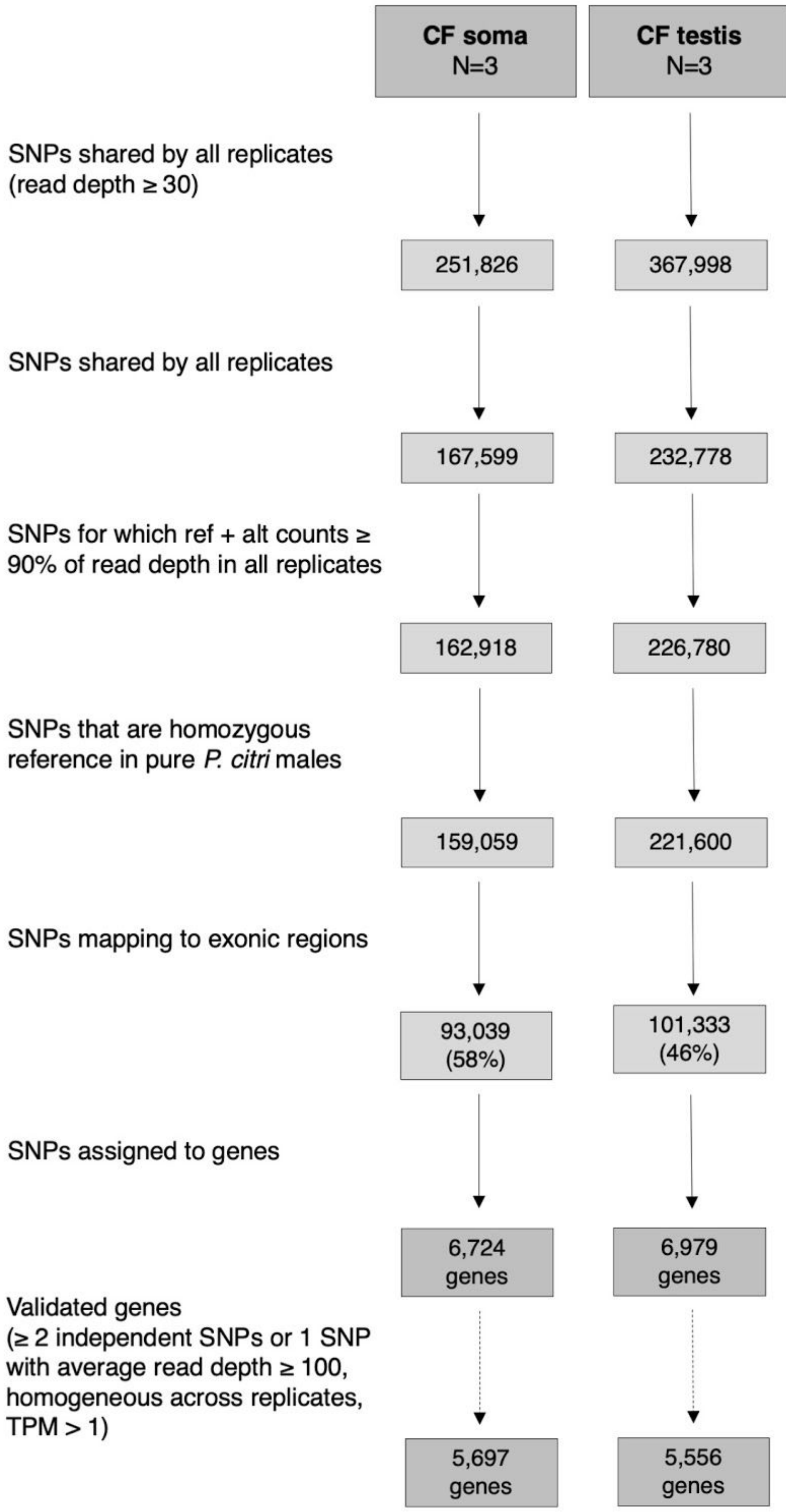


Fig. S1. Workflow for informative SNP filtering and assignment to genes in transcriptomes of CF hybrid males.

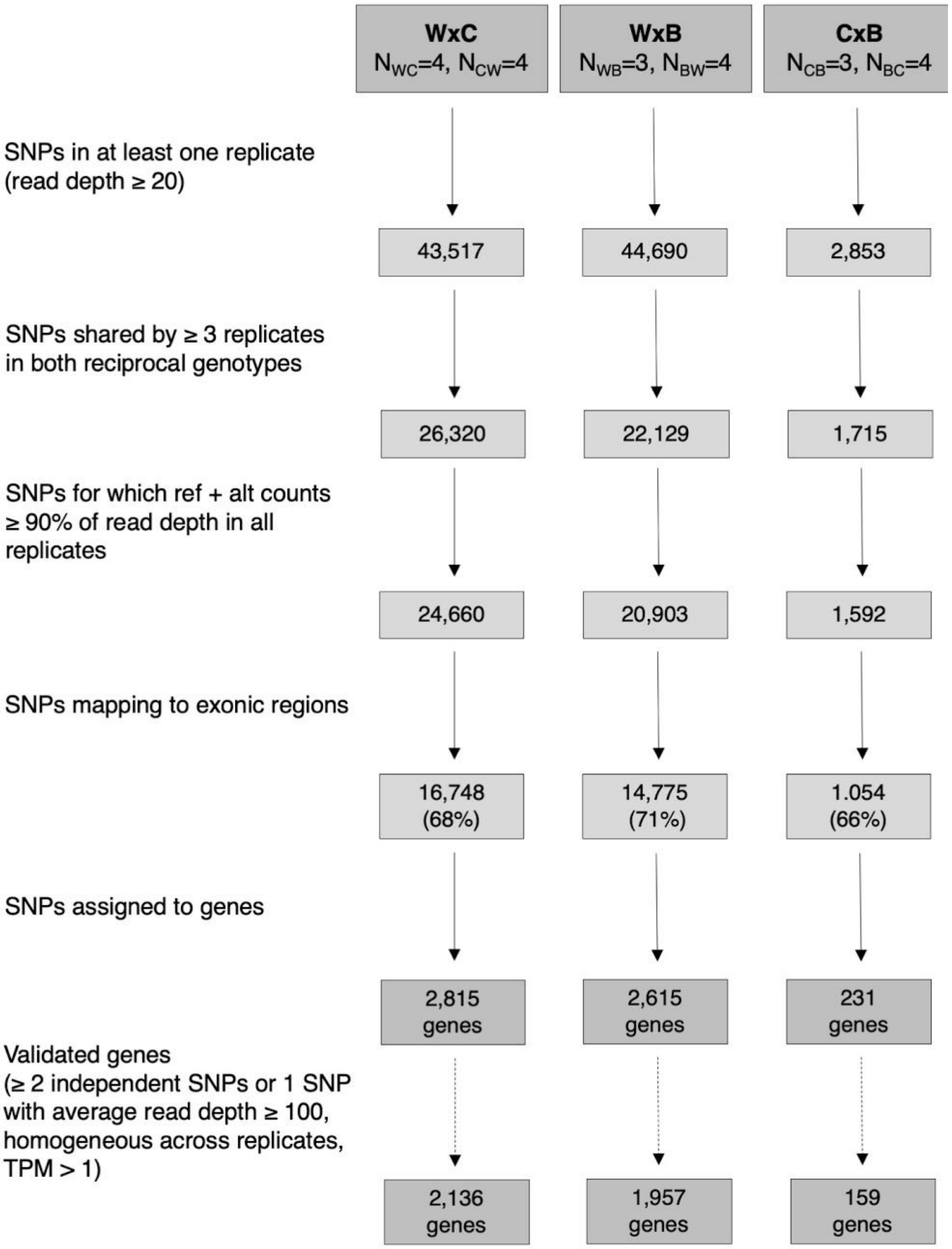


Fig. S2. Workflow for informative SNP filtering and assignment to genes in transcriptomes of intraspecific *P. citri* males

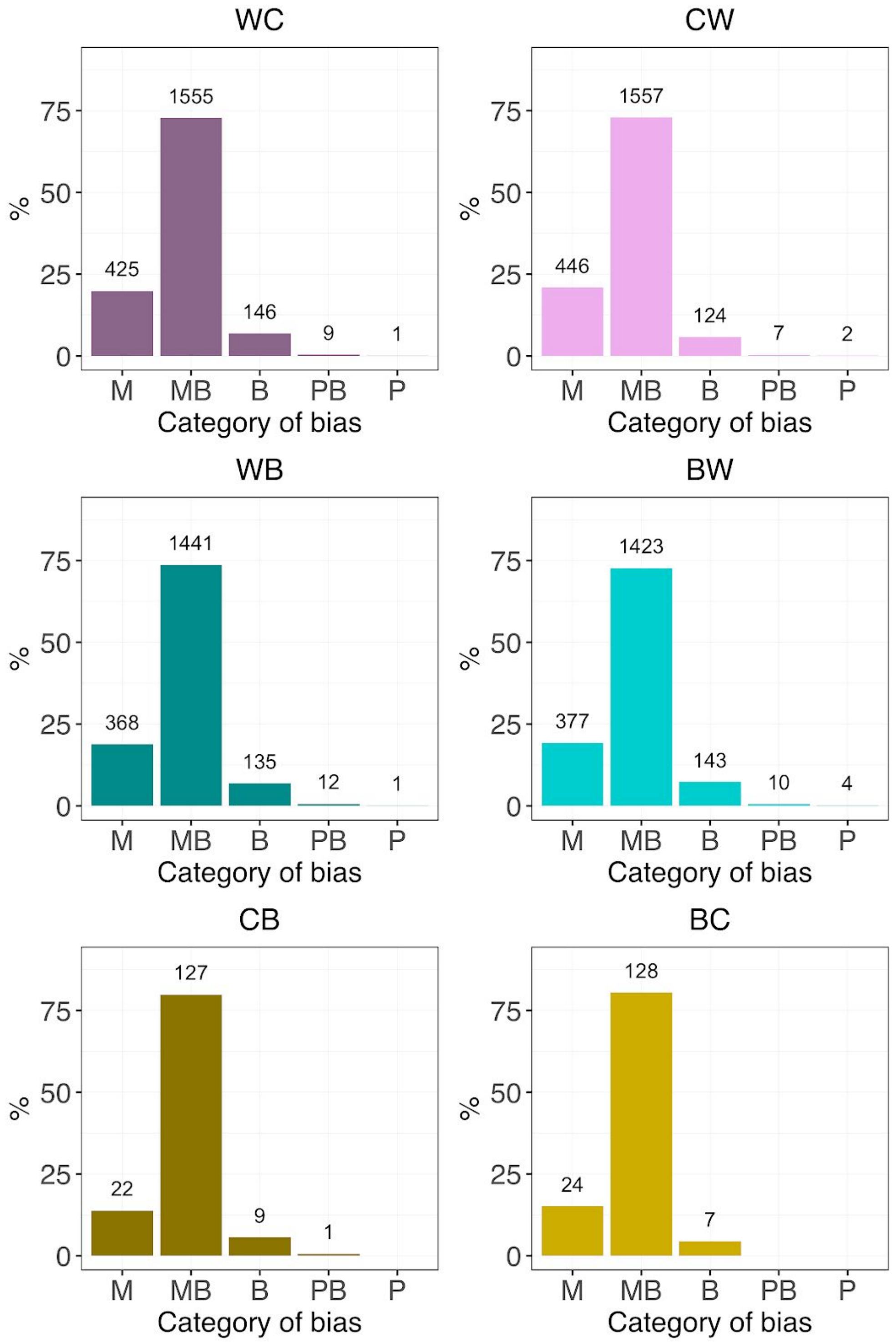


Fig. S3. Counts of genes with allele-specific information in individual intraspecific F1 genotypes according to ASE category (from completely maternal, M, to completely paternal, P)

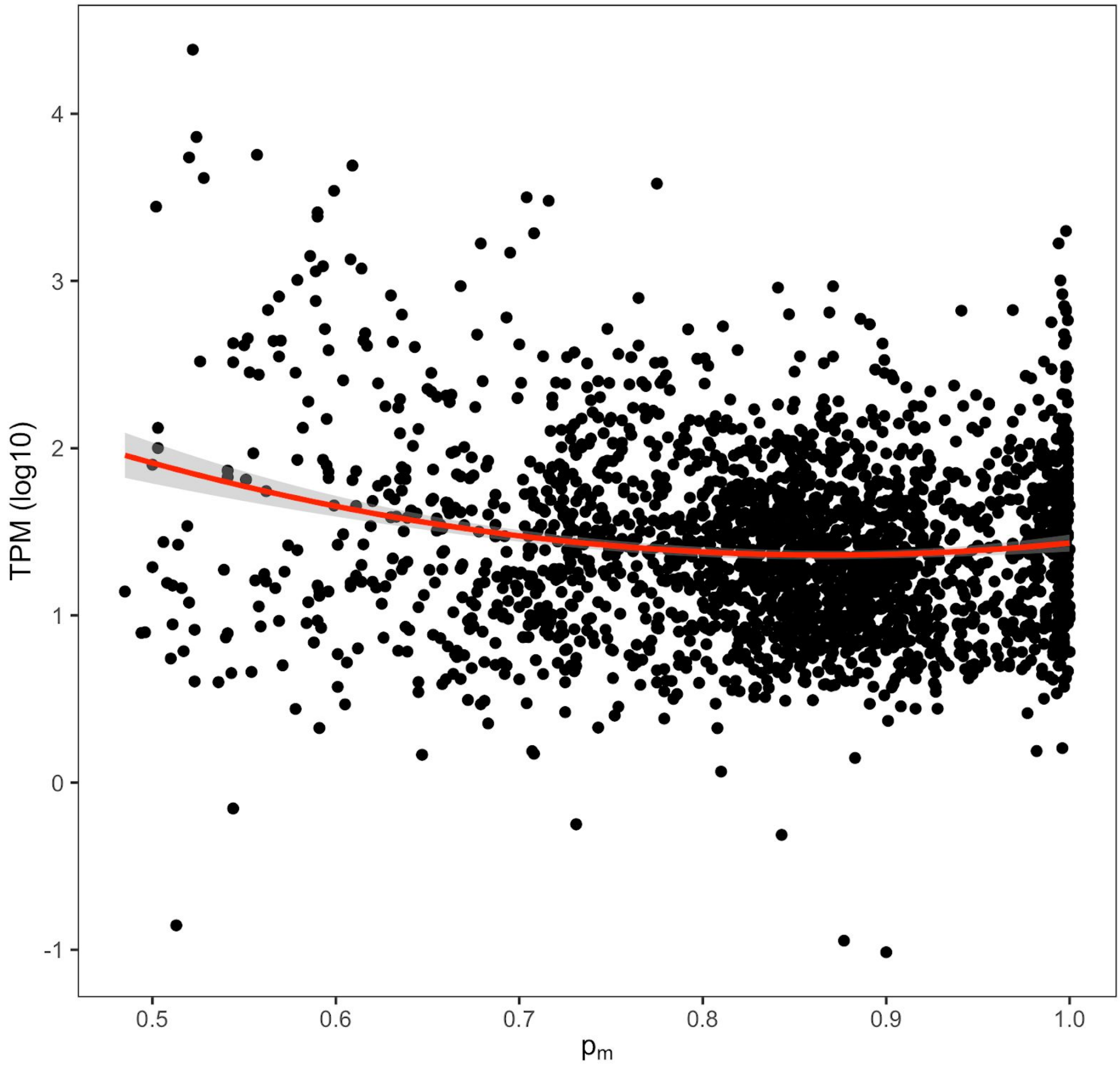


| Term | Estimate | SE | T | P value |
|---|---|---|---|---|
| *Intercept* | 4.4287 | 0.4347 | 10.189 | < 0.0001 |
| $p_m$ | -7.0715 | 1.0937 | -6.465 | < 0.0001 |
| $p_m$^2 | 4.0751 | 5.990 | 5.990 | < 0.0001 |

Fig. S4. Results of the linear model to evaluate the relationship between gene expression levels and bias to the maternal genome (n = 2,379 genes with ASE in intraspecific males).

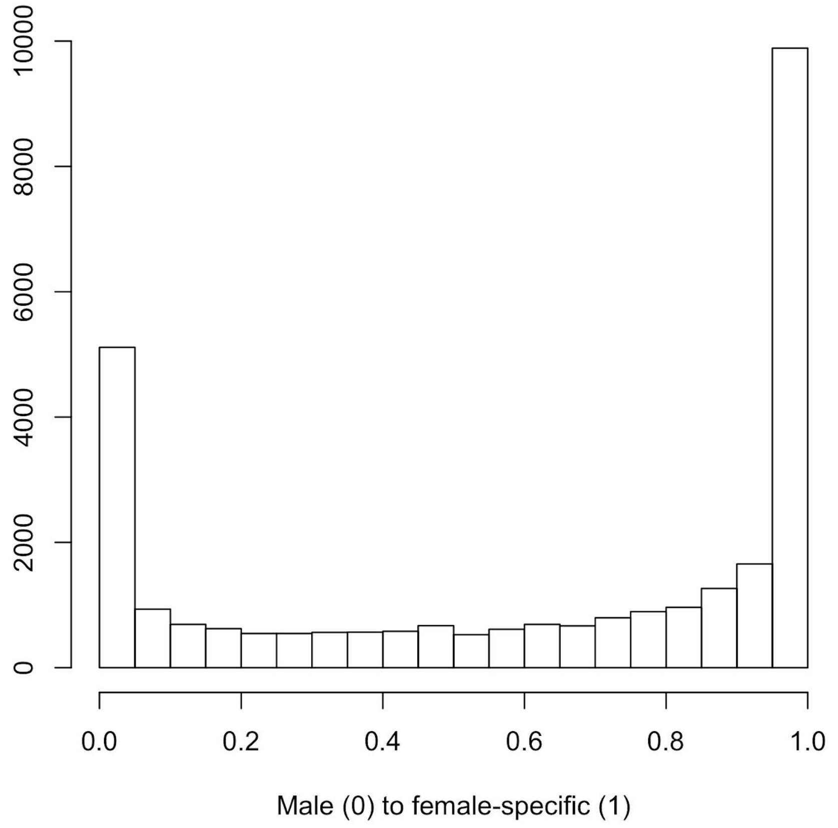


Fig. S5. Distribution of sex-specific expression in *P. citri*

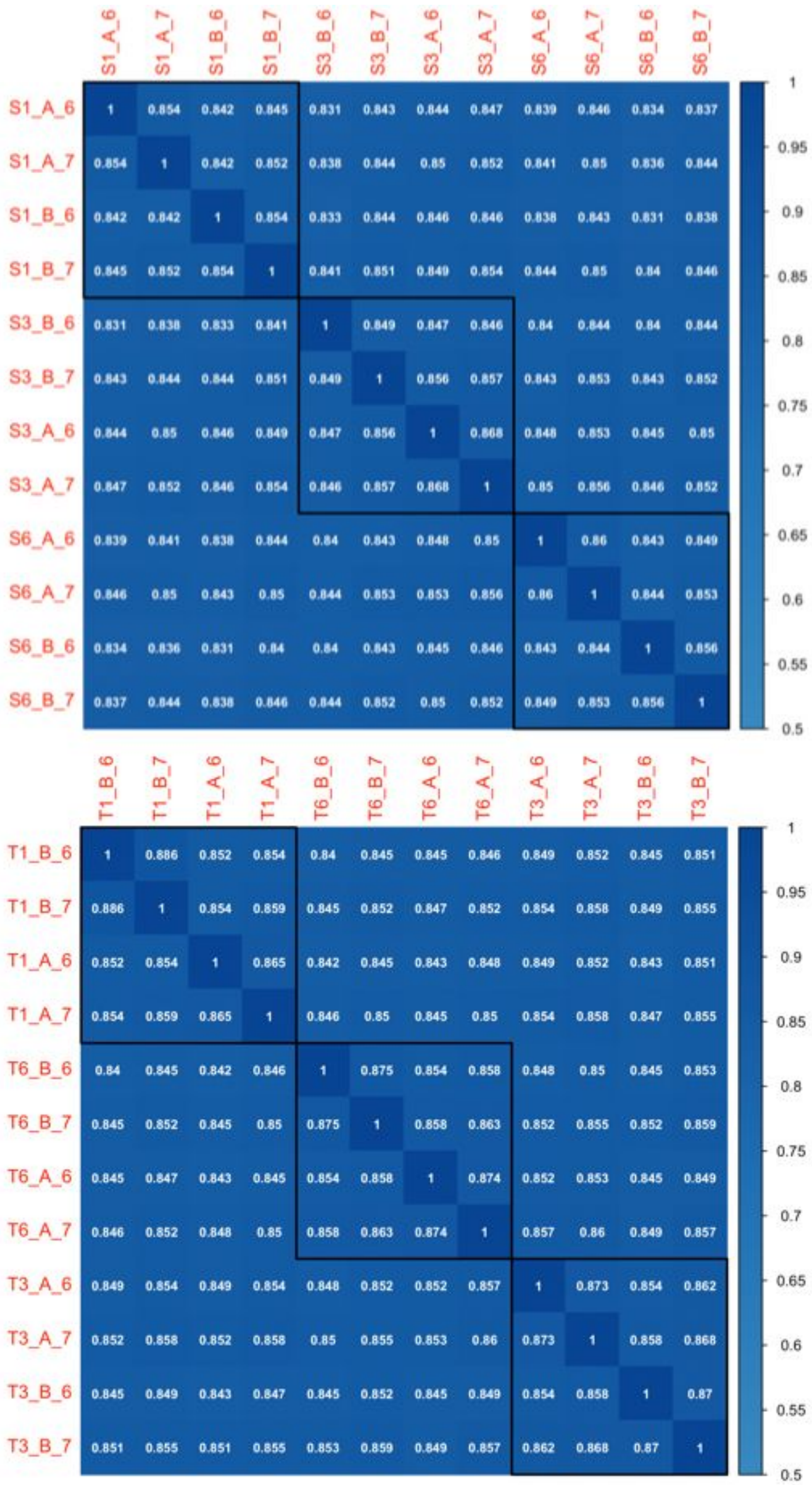


Fig. S6. Correlation matrix heatmaps of gene expression levels (TPM) between samples. Two independent cDNA amplifications (A, B) were performed for each biological replicate of hybrid soma (S1, S3, S6) and hybrid testes (T1, T3, T6). For each tissue, 6 TruSeq Nano libraries were sequenced on two HiSeq 4000 lanes (6, 7). To assess consistency between lane and technical replicates, we mapped each of the 12 soma and testIs RNA-seq datasets separately to the pseudogenome and estimated gene expression values as TPM. The heatmaps show correlation coefficients (Spearman's ρ) across the soma (top panel) and testis (bottom panel) RNA-seq datasets. We used the "hclust" and "addrect" options in the corrplot R package (https://github.com/taiyun/corrplot) to order samples by hierarchical clustering and identify the three main clusters, which correspond to the biological replicates in both hybrid soma and testes.

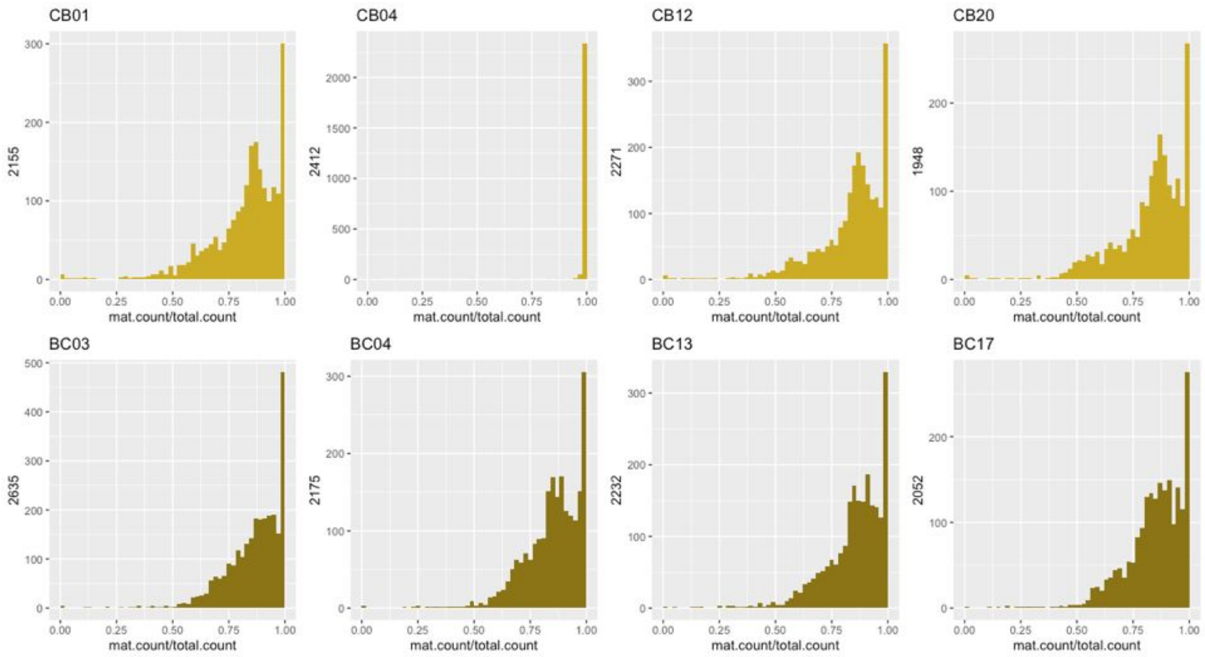


Fig. S7. Histograms of expression biases to maternal genome, $p_m$, at SNP level in all F1 samples from crosses between CP1-2 and BGOX-6 parents (CB, CP1-2 mothers; BC, BC, BGOX-6 parents). CB04 was excluded from the analysis.

Table S1. Counts and average bias to the maternal genome $p_m$ (SD) of validated informative SNPs in CF transcriptomes, grouped by annotation feature.

| ***CF soma*** | **Annotation** | **N SNPs** | **$p_m$** |
|---|---|---|---|
| | Exonic | 93039 (58.5%) | 0.86 (0.13) |
| | Intergenic | 46544 (29.3%) | 0.92 (0.11) |
| | Intronic | 18401 (11.4%) | 0.91 (0.13) |
| | Orphan | 1075 (0.7%) | 0.93 (0.10) |
| ***CF testis*** | **Annotation** | **N SNPs** | **$p_m$** |
| | Exonic | 101333 (45.7%) | 0.96 (0.07) |
| | Intergenic | 81002 (36.6%) | 0.98 (0.06) |
| | Intronic | 36403 (16.4%) | 0.98 (0.05) |
| | Orphan | 2862 (1.3%) | 0.98 (0.08) |

Table S2. Counts and average biases to the maternal genome $p_m$ (SD) within reciprocal genotypes of validated informative SNPs in intraspecific F1 transcriptomes, grouped by annotation feature.

| ***W x C*** | **Annotation** | **N SNPs** | **$p_m$ (WC)** | **$p_m$ (CW)** |
|---|---|---|---|---|
| | Exonic | 16748 (67.9%) | 0.84 (0.13) | 0.85 (0.13) |
| | Intergenic | 6871 (27.9%) | 0.84 (0.15) | 0.86 (0.14) |
| | Intronic | 932 (3.8%) | 0.78 (0.22) | 0.82 (0.18) |
| | Orphan | 109 (0.4%) | 0.85 (0.14) | 0.84 (0.17) |
| ***W x B*** | **Annotation** | **N SNPs** | **$p_m$ (WB)** | **$p_m$ (BW)** |
| | Exonic | 14775 (70.7%) | 0.84 (0.13) | 0.83 (0.15) |
| | Intergenic | 5414 (25.9%) | 0.84 (0.14) | 0.84 (0.16) |
| | Intronic | 603 (2.9%) | 0.80 (0.20) | 0.81 (0.19) |
| | Orphan | 111 (0.5%) | 0.90 (0.12) | 0.87 (0.16) |
| ***C x B*** | **Annotation** | **N SNPs** | **$p_m$ (CB)** | **$p_m$ (BC)** |
| | Exonic | 1054 (66.2%) | 0.81 (0.13) | 0.83 (0.11) |
| | Intergenic | 438 (27.5%) | 0.82 (0.14) | 0.82 (0.15) |
| | Intronic | 53 (3.3%) | 0.84 (0.11) | 0.84 (0.12) |
| | Orphan | 111 (3.0%) | 0.95 (0.11) | 0.96 (0.05) |

Table S3. GO enrichment analysis of 172 genes with biparental (B) or predominantly paternal (PB, P) expression in soma of CF hybrid males against a background population of 3,193 genes with allele-specific information and associated GO terms. All significant GO terms are enriched.

| ***GO term*** | ***Domain*** | ***Ratio*** | ***Ratio in pop*** | ***FDR*** | ***Gene ID*** |
|---|---|---|---|---|---|
| GO:0006412 (translation) | BP | 17/ 172 | 59/ 3193 | 7.7e-5 | g12633, g13997, g1425, g19442, g25371, g26288, g34399, g37033, g38122, g38200, g38206, g38423, g5843, g636, g674, g762, g9816 |
| GO:0055114 (oxidation-reduction process) | BP | 21/ 172 | 171/ 3193 | 4.7e-2 | g10992, g13384, g14250, g14596, g17372, g23111, g2512, g28606, g33094, g34137, g34342, g3639, g36400, g37858, g38672, g39758, g4370, g6152, g7114, g8627, g8706 |
| GO:0005840 (ribosome) | CC | 17/ 172 | 58/ 3193 | 2.1e-5 | g12633, g13997, g1425, g19442, g25371, g26288, g34399, g37033, g38122, g38200, g38206, g38423, g5843, g636, g674, g762, g9816 |
| GO:0003735 (structural constituent of ribosome) | MF | 17/ 172 | 60/ 3193 | 1.3e-4 | g12633, g13997, g1425, g19442, g25371, g26288, g34399, g37033, g38122, g38200, g38206, g38423, g5843, g636, g674, g762, g9816 |
